# Contribution of non-coding mutations to *RPGRIP1*-mediated inherited retinal degeneration

**DOI:** 10.1101/211292

**Authors:** Farzad Jamshidi, Emily M. Place, Sudeep Mehrotra, Daniel Navarro-Gomez, Mathew Maher, Elise Valkanas, Timothy J. Cherry, Monkol Lek, Daniel MacArthur, Eric A. Pierce, Kinga M. Bujakowska

**Affiliations:** Ocular Genomics Institute, Department of Ophthalmology, Massachusetts Eye and Ear Infirmary, Harvard Medical School, Boston, MA; Program in Medical and Population Genetics, Broad Institute of MIT and Harvard, Cambridge, MA; Center for Developmental Biology and Regenerative Medicine, Seattle Children’s Research Institute and University of Washington, Department of Pediatrics, Seattle, WA; 4Analytic and Translational Genetics Unit, Massachusetts General Hospital, Boston, MA

**Keywords:** inherited retinal degeneration, non-coding mutations, *RPGRIP1*, intronic mutations, whole-genome sequencing

## Abstract

**Purpose:** With the advent of gene therapies for inherited retinal degenerations (IRDs), genetic diagnostics will have an increasing role in clinical decision-making. Yet the genetic cause of disease cannot be identified using exon-based sequencing for a significant portion of patients. We hypothesized that non-coding mutations contribute significantly to the genetic causality of IRDs and evaluated patients with single coding mutations in *RPGRIP1* to test this hypothesis.

**Methods:** IRD families underwent targeted panel sequencing. Unsolved cases were explored by whole exome and genome sequencing looking for additional mutations. Candidate mutations were then validated by Sanger sequencing, quantitative PCR, and *in vitro* splicing assays in two cell lines analyzed through amplicon sequencing.

**Results:** Among 1722 families, three had biallelic loss of function mutations in *RPGRIP1* while seven had a single disruptive coding mutation. Whole exome and genome sequencing revealed potential non-coding mutations in these seven families. In six, the non-coding mutations were shown to lead to loss of function *in vitro*.

**Conclusion:** Non-coding mutations were identified in 6 of 7 families with single coding mutations in *RPGRIP1*. The results suggest that non-coding mutations contribute significantly to the genetic causality of IRDs and *RPGRIP1*–mediated IRDs are more common than previously thought.

## INTRODUCTION

Inherited retinal degenerations (IRDs) are a group of monogenic diseases that are the most common cause of blindness in the working age population^1^. About 260 genes have been associated with IRDs with functions spanning almost every aspect of cellular function: from splicing machinery, to microtubular transport and phototransduction^1^. State-of-the-art clinical diagnostics using next-generation sequencing (NGS) of known IRD genes successfully identifies the causal mutation in only 50 to 70% of cases^2,3^. Although copy number changes^4^ and intronic mutations^5^ contribute to disease, they are not routinely assessed and likely contribute to the genetic causality in a significant portion of currently unsolved cases. With the advent of successful gene therapies for IRDs^6^, understanding such non-coding mutations and developing assays to evaluate them is of increasing importance. One example is the autosomal recessive *RPGRIP1*-associated disease which is an attractive candidate for gene therapy with already established success in murine^7^ and canine^8^ models. Yet, almost all of the mutations in *RPGRIP1* have been described in the coding-region^9^.

RPGRIP1 plays a critical role in opsin trafficking, outer-segment disc organization and photoreceptor survival^10,11^. While it primarily localizes to the transition zone of rods and cones, various of its isoforms can be found in the outer segment, along the microtubules as well as in the amacrine cells of the inner plexiform layer^12,13^. Its largest transcript variant, NM_020366, is composed of 3861 coding base pairs distributed over 24 exons^14^. This encodes a 1287 amino acid protein that interacts with a variety of other IRD proteins such as Retinitis Pigmentosa GTPase Regulator (RPGR), SPATA7 and NPHP4^15^. The expression of *RPGRIP1* is limited to the retina and testis^16^.

Confident genetic diagnosis with *RPGRIP1* as the causal gene will be crucial for effective clinical trials of potential therapies. In our analysis of IRD families with targeted panel sequencing of coding regions of IRD-associated genes^17^, we repeatedly noted identification of single likely pathogenic variants in *RPGRIP1* in families without mutations in other IRD disease genes. To test the hypothesis that mutations in *RPGRIP1* are the likely cause of disease in these families, we performed whole exome (WES) and genome sequencing (WGS) to search for non-coding mutations and structural variations accounting for the loss of function (LoF) of the second allele.

## MATERIALS AND METHODS

### Ethical guidelines

The study was approved by the institutional review board at the Massachusetts Eye and Ear (Human Studies Committee MEE in USA) and adhered to the Declaration of Helsinki. Informed consent was obtained from all individuals on whom genetic testing and further molecular evaluations were performed.

### Clinical evaluation

All the patients in this study underwent clinical assessment by ophthalmologists sub-specializing in inherited retinal degenerations. The clinical characteristics are outlined in Table 1.

**Table 1.** Clinical characteristics and mutations in RPGRIP1-mediated inherited retinal degeneration patients

| ID<sup>a</sup> | Gender | Age at Dx | Variant 1<sup>b</sup> | MAF<sup>c</sup> | Variant 2<sup>b</sup> | MAF<sup>c</sup> | Signs<sup>d</sup> | Method |
| --- | --- | --- | --- | --- | --- | --- | --- | --- |
| 237-523 | F | Early | c.3793ins4 | 4×10<sup>-6</sup> | Exon1-2 dup | N/A | Keratoconus, PSC, | GEDi |
|  |  | childhood | (p.V1265Gfs*19) |  |  |  | asteroid hyalosis, ↓ | WES |
|  |  |  |  |  |  |  | CV | WGS |
| 281-608 | M | Early | c.1615_1624del10 | 4×10<sup>-6</sup> | Exon2 dup | N/A | Nystagmus, ON | GEDi |
|  |  | childhood | (p.E539Qfs*2) |  |  |  | atrophy, ↓ CV | WGS |
| 601-1236 | M | Infancy | c.3238+1G>A | 0 | c.1611+27G>A | 0 | Nystagmus, scatter | GEDi |
|  |  |  |  |  |  |  | hypopigmented spots | WGS |
|  |  |  |  |  |  |  | in retinal periphery |  |
| 949-1907 | M | Infancy | c.3618-1_3621del5 | 0 | c.1468-263G>C | 0 | Nystagmus, | GEDi |
|  |  |  |  |  |  |  | peripheral atrophy, | WGS |
|  |  |  |  |  |  |  | pigmentary macular changes |  |
| 827-1591 | M | 1.5 yo | c.895_896del2 | 0 | c.2367+23delG | 2.3×10<sup>-4</sup> | Nystagmus, macular | GEDi |
|  |  |  | (p.G299Sfs*21) |  |  |  | atrophy | WES |
| 1797-3128 | M | 15 yo | c.2302C>T<br>(p.R768*) | 2×10<sup>-5</sup> | Exon19 del | N/A |  | GEDi |
| 79-194 | F | Infancy | c.3793ins4<br>(p.V1265Gfs*19) | 4×10<sup>-6</sup> | c.3793ins4<br>(p.V1265Gfs*19) | 4×10<sup>-6</sup> | Peripheral<br>pigmentary changes,<br>bull's eye macular<br>changes | GEDi |
| 501-336 | F | 4 months | c.2302C>T<br>(p.R768*) | 2×10<sup>-5</sup> | c.711_711del1<br>(p. P237Pfs*40) | 0 | Nystagmus | GEDi |
| 690-1378 | F | Infancy | c.1084_1087del<br>(p.E362NAfs*12) | 0 | c.767C>G<br>p.S256* | 0 | Nystagmus | GEDi |

Dup, duplication; del, deletion; PSC, posterior subcapsular cataract; CV, color vision; ON, optic nerve; N/A, not available; yo, years old.
<sup>a</sup> IDs are presented as Ocular Genomics Institute (OGI) family numbers followed by the individual patient number.
<sup>b</sup> RPGRIP1 mutations in each of two alleles, with protein alteration in parenthesis
<sup>c</sup> Minor allele frequency based on the Genome Aggregation Database (gnomAD)
<sup>d</sup> All patients exhibited diminished visual field, visual acuity, and electroretinography signals in addition to attenuated vessels and bone spicules on fundoscopy. Additional finding are indicated in the table.

### Sequencing

DNA was extracted from venous blood using the DNeasy Blood and Tissue Kit (Qiagen, Hilden, Germany). All samples underwent GEDi sequencing as described previously^17^. Whole exome and genome sequencing were done at the Center for Mendelian Genomics at the Broad Institute of MIT and Harvard using methodology described previously^18^. Sanger sequencing was performed on ABI 3730xl using BigDye Terminator v3.1 kits (Life Technologies, Carlsbad, CA) and using PCR primers indicated in Supplementary Information. When PCR products were sequenced, they were purified prior to sequencing (ExoSap-IT, Affymetrix, Santa Clara, CA). Gel bands that were Sanger sequenced had DNA extracted via the Zymoclean^TM^ Gel DNA Recovery Kit (Zymo Research, Irvine, CA).

### Bioinformatics

Analyses of DNA sequence data were performed as described previously^17,19^. Briefly, BWA was used for alignment. SAMtools and custom programs were used for single nucleotide polymorphism and insertion/deletion calls^19^. Variants of interest were limited to polymorphisms with less than 0.005 allelic frequency in the gnomAD (http://gnomad.broadinstitute.org/) and ExAC (http://exac.broadinstitute.org/) databases^18^. Whole genome copy number analysis, with consideration of structural changes, was done using Genome STRiP 2.0^20^. For the analysis of splicing patterns from amplicon-sequencing, STAR (version 2.5.3a) aligner^21^ was used to generate an index of the human genome (GRCh37.75.dna.primary_assembly.fa) and to align the reads. IGV^22^ was used to load the aligned sequences (BAM files) and for data visualization with Sashimi plots.

### Polymerase chain reactions (PCR), cloning and site-directed mutagenesis

PCR was performed using PfuUltra II Fusion polymerase (Agilent Technologies, Santa Clara, CA) on genomic DNA of patients harboring the mutations of interest (primers are listed in the Supplementary). The PCR products were cloned into pENTR Directional TOPO vector (Thermo Fisher, Waltham, MA) and used to transform chemically competent *Escherichia coli* (One Shot TOP10, Thermo Fisher, Waltham, MA). Plasmid DNA from single colonies was extracted with miniprep kits (ZymoPURE, Zymo Research) and analyzed by restriction enzyme digestion with BsrGI (NE Biolabs, Ipswich, MA) and Sanger sequencing. Essential splice-site mutations were introduced by site-directed mutagenesis (QuickChange II Site Directed mutagenesis kit, Agilent Technologies) and verified by Sanger sequencing. Colonies with the correct sequence and restriction enzyme pattern were then sub-cloned into the pCS2+GW vector (kind gift from Dr Erica Davis) via Gateway LR clonase II (Thermo Fisher) and similar analyses as before was done to isolate vectors with the appropriate inserts for transfection experiments. The final vector included *RPGRIP1* exons 11-16 including extensions into intron 10 and 16 on the 5’ and 3’ ends, which was cloned into pCS2+GW and used for splicing assays.

### Quantitative polymerase chain reactions (qPCR)

Five□nanograms (ng) of genomic DNA, 200 nM of each primer and 10 µl of Fast SYBR Green Master Mix (Life Technologies, Grand Island, NY) were used for qPCR reactions which were performed on a Stratagene Mx3000P instrument (Agilent Technologies) using the standard thermocycling program (95 °C for 3□min, 40 cycles of 95 °C for 20□s, and 60 °C for 1□min, followed by a melting curve). The ddCT method was used for the analysis of results where *ZNF80* was used as a reference gene and an in-house DNA sample with wild type *RPGRIP1* (OGI200) used for normalization. Each sample was tested in triplicate and the average value was used. Standard deviation with error propagation was used to calculate up and down errors.

### Cell culture and transfections

Human embryonic kidney (HEK293T) and retinoblastoma (WERI-Rb1) cells purchased from American Type Culture Collection (ATCC, Manassas, VA) and maintained in RPMI medium supplemented with 10% Fetal Bovine Serum-1640 (Thermo Fisher). 2ml of 5 x 10^5^ cells/ml were plated into each well of a 6-well plate (Corning Inc., Corning, NY) 12hrs prior to transfections. 1-5 μg of vector DNA per well was used for transfections using a commercial reagent (Lipofectamine 2000, Invitrogen, Carlsbad, CA). Cells were harvested for RNA extraction 48 hours after transfection.

### RNA isolation and cDNA synthesis

Cells were lysed with TRIzol (Thermo Fisher). After BCP or chloroform (Sigma-Aldrich, St. Louis, MO) treatment, the aqueous phase was transferred to mRNeasy columns with DNase I digestion performed on-column (Qiagen). Quantification was performed via NanoDrop (Thermo Fisher) and 500ng of RNA was converted to cDNA using oligo(dT) primers and SuperScript II (Thermo Fisher).

### Splicing assay and amplicon sequencing

The mutant, control and wild-type vectors described above were transfected in to HEK293T and WERI-Rb1 cells. Two days post-transfection, cDNA was generated as described and RT-PCR performed amplifying the flanking exons of the point mutation of interest. The PCR products were purified by DNA-clean and concentrator kits (Zymo Research). Amplicon sequencing was then performed at the Massachusetts General Hospital Center for Computational and Integrative Biology, where the PCR products were fragmented using sonication and sequenced with the standard NGS pipeline. Visualization and analysis of the data were performed as described under bioinformatics.

### Epigenetic features surrounding the RPGRIP1 locus

ATAC-Seq and ChIP-Seq were performed according to previously published methods^23,24^. Briefly, human retinal tissue was obtained from donors 25-65 years old with a postmortem interval <8hrs (Lions Vision Gift, Portland, OR). Approximately 20,000 nuclei were isolated for ATAC-Seq and the transposition reaction was performed for 60min at 37C. ChIP-Seq was performed on approximately 25 million cells per reaction using the following antibodies: CTCF (Abcam AB70303); H3K4me2 (Abcam AB7766 lot GF160184-1; RRID AB_2560996); H3K27ac (Abcam AB4729, lot GF150367-1; RRID AB_2118291); CRX (Santa Cruz, B11X, lot E1409); OTX2 (Abcam, AB21990, lot GR242019-1); NRL (Abcam, AB137193, lot GR104520-2); RORB (Diagenode, pAb-001-100, lot HS-0010); MEF2D (Greenberg lab, 2373). All sequencing was performed on an Illumina NextSeq500 to a depth of >10M reads. Mapping, alignment and normalization of reads was peformed as previously described^24^. Genome tracks were displayed using the UCSC genome browser.

## RESULTS

Genetic analysis of 1722 IRD probands that underwent targeted exon sequencing of known IRD genes^17^ revealed three patients with bi-allelic loss-of-function (LoF) mutations in *RPGRIP1* and seven with only one LoF change in this gene (Table 1, Figure S1). In the latter seven families no other significant mutations in *RPGRIP1* or other IRD genes were identified. Since they were all diagnosed with an early onset IRD, a characteristic presentation of *RPGRIP1* disease^25^, further testing was performed to search for non-coding and structural variants in *RPGRIP1* or other IRD genes through whole exome and whole genome sequencing. Among these seven families, three of the second-allele mutations were predicted to be copy number changes and three were intronic mutations (Table 1). In one family, OGI-578, we did not validate second pathogenic variants in *RPGRIP1* (Figure S1).

WES and WGS studies showed that three families have structural variants as the second mutation in *RPGRIP1*. Analysis of WES data showed that patient OGI-1797-3128 had a predicted deletion of exon 19 (Table 1), which was confirmed by qPCR (Figure S2). Exon 19 has 139 nucleotides, and thus its deletion is predicted to lead to a frameshift resulting in a premature stop codon and likely subsequent nonsense mediated decay (NMD)^26^. In OGI-281-608, the gain of a copy of exon 2 was detected through structural analysis^20^ in addition to coverage-based predictions of WGS data. The structural change led to misalignment of paired-end reads (Figure 1c and 1d), which indicated a tandem duplication. This mutation was validated by qPCR (Figure 1b), and the predicted breakpoint confirmed through PCR and Sanger sequencing (Figure 1e and 1f). Sanger sequencing also identified 135bp of missing DNA upstream of the breakpoint, suggesting a possible complex rearrangement as the causal event^27^. Given that exon 2 is 133bp, its tandem duplication would lead to a LoF allele.

**Figure 1.**
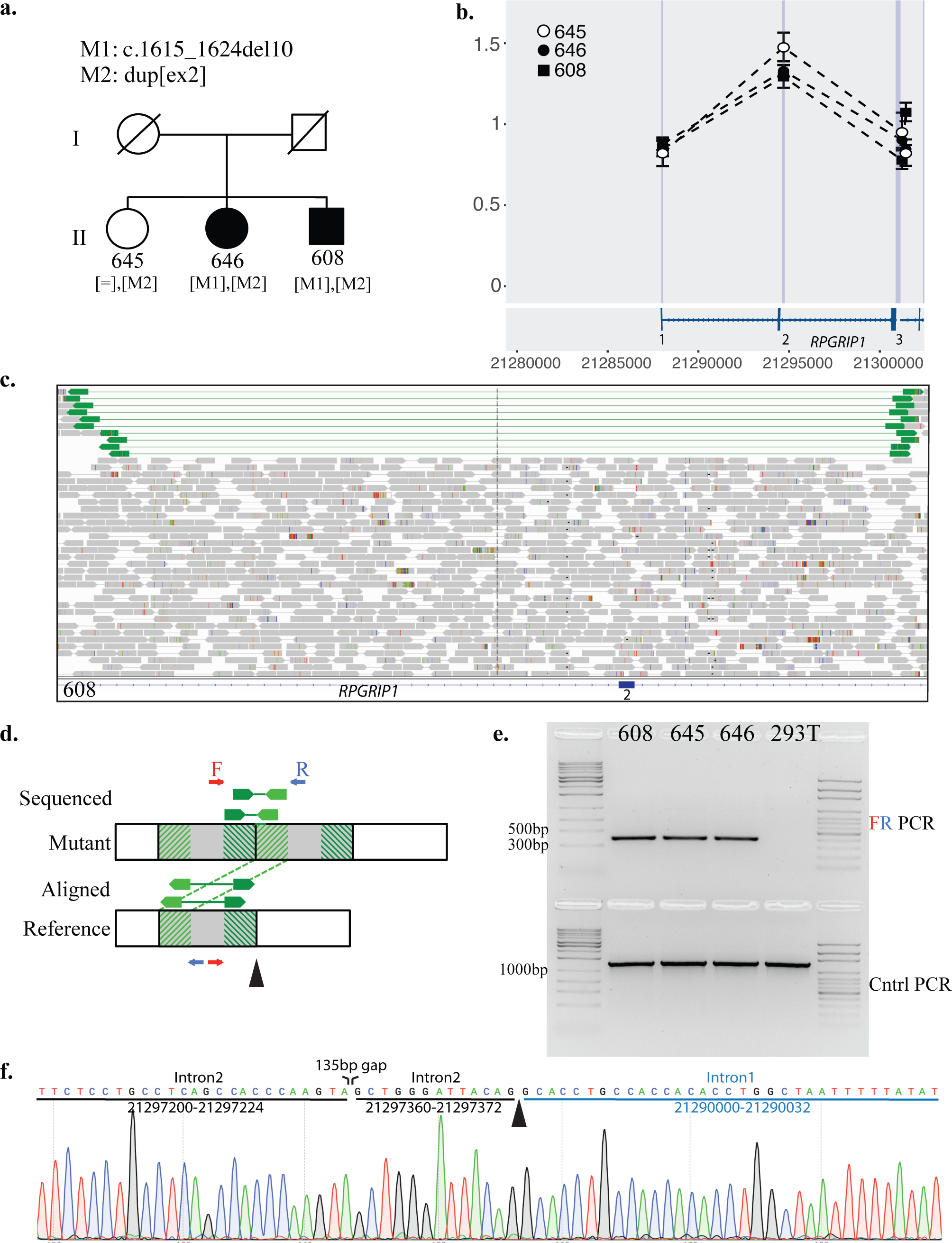
Exon 2 duplication in OGI-281. **(a)** Pedigree of the family showing deceased parents and the three siblings all of whom were analyzed. **(b)** qPCR based copy number results along the first three exons of *RPGRIP1*. All three siblings have a duplication of exon 2 in an *RPGRIP1* allele. Exons 1 and 3 are not affected. The bottom panel shows the locations of *RPGRIP1* exons based on the NM_020366 transcript. **(c)** Integrative genomics viewer^22^ (IGV) view of the sequenced whole genome sequencing (WGS) reads where the duplication was discovered for OGI-281-608. The bottom of the figure shows the location of exon 2 of *RPGRIP1.* The gray thick arrows correspond to expected paired-end reads. The green thick arrows are mapped reads that have aligned abnormally and hint to a mutation. **(d)** Schematic explanation of the how genomic duplication would lead to the abnormal paired-end reads seen in Figure 1c. The gray region is the area of hypothetical duplication, while the green thick arrows are the paired-end reads that will align abnormally. The top of the figure shows what is actually sequenced in the mutant sample, while the bottom shows how such sequenced reads would map to a reference wild-type (WT) model. The aligned pared-end reads of the mutant will have a greater distance between them and will point away form one another as seen in Figure 1c. The dark and light green hash lines correspond to the aligning sequenced of the paired-end reads. The primers used for Figure 1e are shown as blue and red arrows. Their directionality is indicated relative to the mutant (top) and WT models (bottom). **(e)** Polymerase chain reaction (PCR) across the predicted duplication breakpoint using primers represented in Figure 1d. Presence of a tandem duplication would yield a product while its absence would lead to no amplification as the primers would be pointing away from each other. The predicted duplication is present in all OGI-281 family members while it is absent in HEK293T cells. The control (Cntrl) PCR on the bottom was done to ensure that larger products could be amplified from all samples thus ensuring DNA fragmentation or quality was not a confounding factor. **(f)** Sanger sequencing identifying the exact breakpoint (black arrowhead) using OGI-281-608 PCR product from (e). There is a 135bp deletion, upstream of the breakpoint.

Patient OGI-237-523 similarly had a tandem copy number gain but in both exons 1 and 2 detected by WES (Figure 2b). The 5’ breakpoint was mapped 2Kb upstream of exon 1 (Figures 2c, S3). However, given that the second copy would maintain 2Kb upstream of the exon 1, which would include a proximal presumed promoter (Figure 2c), we questioned whether a second transcriptional start site (TSS) within the duplicated 5’ upstream region would exclude the mutant exons 1’ and 2’ thus leading to normal transcription. Therefore, we hypothesized that perhaps a critical *RPGRIP1* regulatory domain exists outside of 2Kb region upstream of exon 1. Review of transcriptome data of normal human retina^28^ revealed an additional exon upstream of the annotated transcript, which was identified by numerous split reads as well as high sequence coverage approximately 8Kb upstream of the annotated *RPGRIP1* exon 1 (Figure 2d). We further assessed this using ATAC-seq and ChIP-seq data, which showed open chromatin and transcription factor binding to this region and not to the previously annotated *RPGRIP1* TSS (Figure 2E). These results suggest that the retinal TSS of *RPGRIP1* and its regulatory regions are 8Kb upstream of the currently annotated exon 1. This also explains the likely pathogenic effect of the structural mutation seen in OGI-237-523. The tandem exon 1 and 2 duplication excludes a TSS and will lead to a premature stop-codon 29 codons past the start of the duplicated 5’UTR of exon 1.

**Figure 2.**
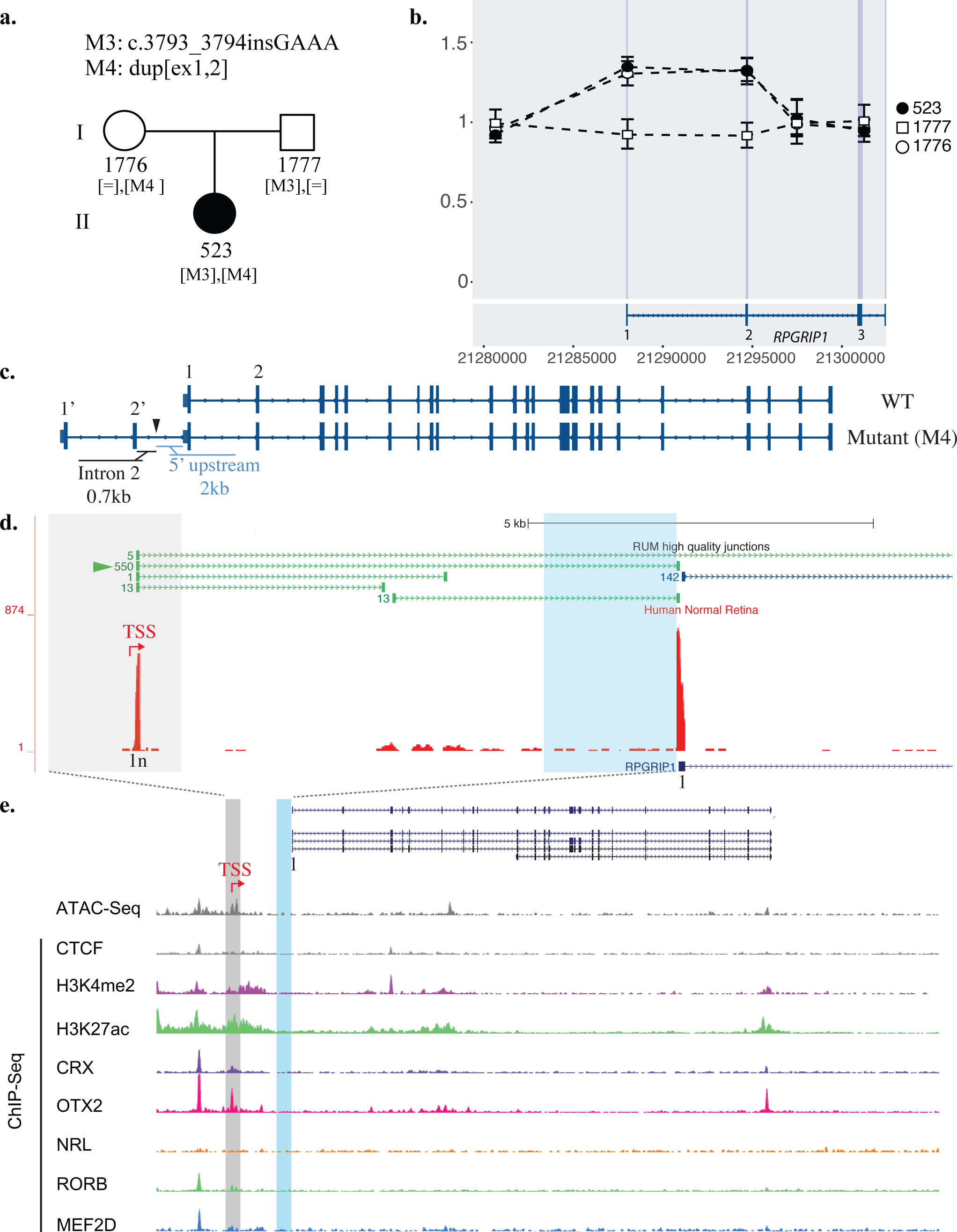
Exon 1 and 2 duplication in OGI-237 and identification of a novel exon. **(a)** Pedigree of OGI-237 showing the segregation of the *RPGRIP1* mutations in the family. **(b)** qPCR of OGI-237 family members, showing the presence of duplication of both exons 1 and 2 in the mother and the proband (523) while the father has a normal copy number across these exons. The bottom panel shows the locations of *RPGRIP1* exons based on the NM_020366 transcript. **(c)** Representation of the predicted WT and mutant alleles with the duplication of exons 1 and 2 (M4). Arrowed lines are introns, tall bars are exons, and short bars indicate untranslated regions (UTR). Further analysis (Figure S3), shows that M4 is a result of tandem duplication with a breakpoint at ~2Kb upstream of exon 1 (black arrow-head). If the transcriptional start site (TSS) is within this 2Kb region, then M4 could lead to normal transcripts given uninterrupted exon 1 and its upstream region. **(d)** Exploration of the retina transcriptome^28^ shows however, that there is an additional novel exon (1n) upstream of the currently annotated exon 1 in the canonical NM_020366 transcript model. Thus the actual TSS is expected to be upstream of exon 1n rather than exon 1. The red bars are indicative of read depth of the transcriptome data. The green and blue arrowed lines at the top indicate split reads between exons. The green are across unannotated exons, while the blue corresponds to annotated exons. There are 550 split reads between exons 1n and 1, further confirming the presence of this novel exon. The NM_020366 canonical transcript model is shown at the bottom. The gray and light blue highlighted areas corresponded to the gray and light blue areas of Figure 2e respectively. **(e)** ATAC-Seq and ChIP-Seq from adult human retina of histone modifications and transcription factor binding at the *RPGRIP1* locus as in (d). The light blue shading represents the area directly upstream of the annotated RPGRIP1 transcript. The light gray shading represents a promoter region and newly-discovered retina-specific exon suggested by RNA-Seq, ATAC-Seq and ChIP-Seq. The transcript models of *RPGRIP1* are shown on top.

Three families with single mutations identified in *RPGRIP1* by panel-based genetic testing had deep intronic mutations that segregated in the families and had low frequencies in the gnomAD database^18^ (Table 1). We hypothesized that these mutations could potentially lead to aberrant splicing creating LoF alleles as well. To assess the effect of these changes on splicing, midi-gene assays were performed in HEK293T cells with mutant and wild-type (WT) constructs, as well as essential splice-site changes serving as positive controls. In OGI-827-1591, the c. 2367+23delG change led to a significant retention of intron 15 in mutant vs. wildtype comparable to the findings in the c. 2367+1G>A positive control (Figure 3a). While the ratio of splicing between exons 15 to 16 to that between exons 14 and 15 was 6090/6207 or 0.98 in wild-type, this proportion was altered to 2209/3982 or 0.55 in mutant. This suggests a relative reduction of 43% of the appropriate splicing at this locus. The aberrant splice isoform leads to a premature stop-codon 19 codons into intron 15.

**Figure 3.**
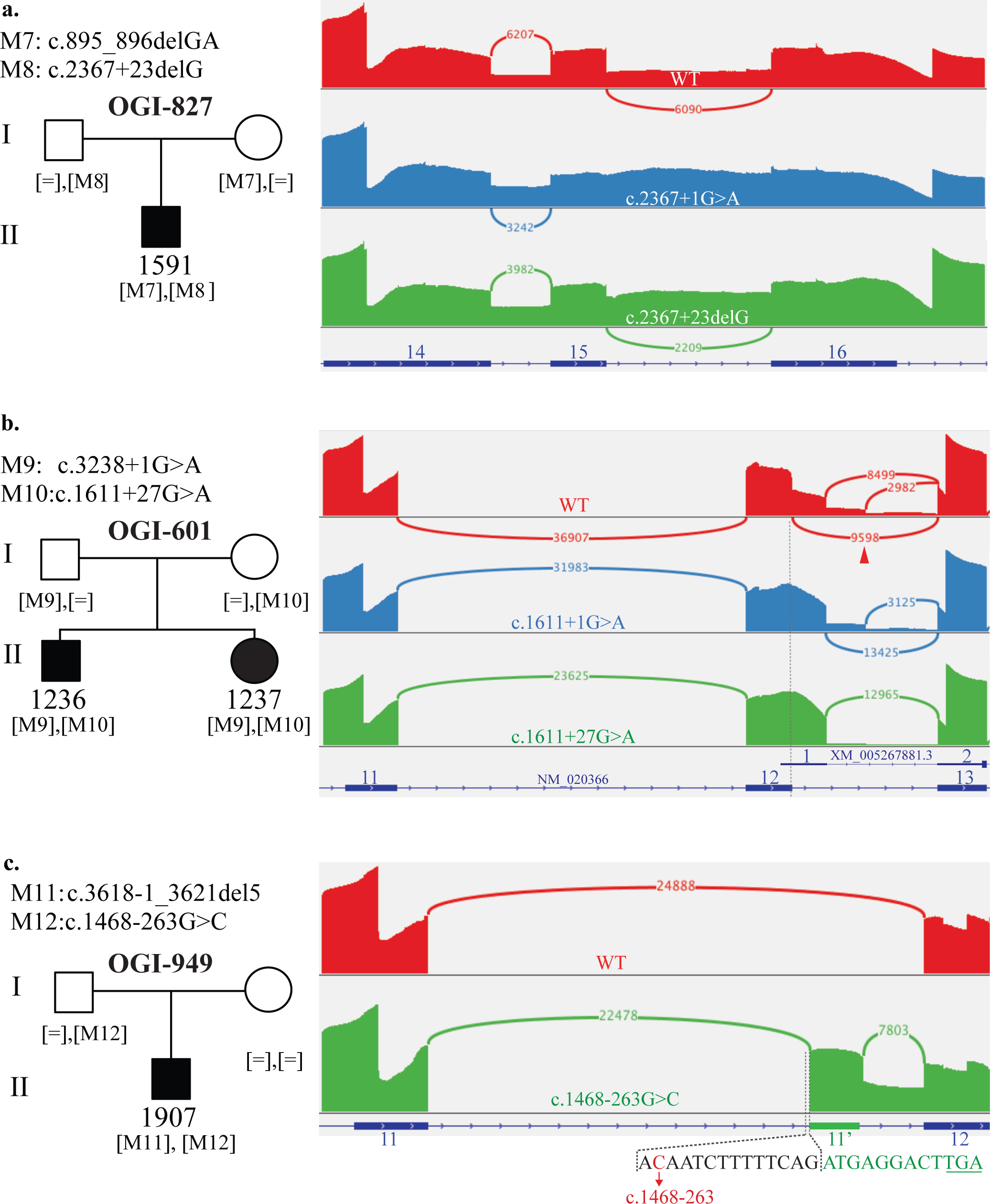
Sashimi plots of the amplicon sequencing of splicing assays. The sashimi plots are generated using amplicon sequencing on RT-PCR products across the indicated regions in HEK293T cells that were transfected with WT (red), positive control (blue), or mutant (green) *RPGRIP1* vectors. The annotated NM_020366 exons are indicated on the bottom of each plot in dark blue. **(a)** In OGI-827, the intronic mutation caused increased intron retention (green), similar to the essential splice site mutation control (blue). **(b)** In OGI-601, as described in the text, the intronic mutation shifts splice to extend exon 12 (green), as does the essential splice site mutation control (blue). Interestingly, the transcript variant XM_005267881.3 has been reported to include this extention in its exon 1 which is entirely untranslated. Untranslated and translated regions of the transcript models are indicated as thin and thick dark blue lines respectively. **(c)** In OGI-949, the deep intronic mutation leads to inclusion of a cryptic exon 11’ in the mutant transcript (green). The dashed lines highlight the immediate upstream sequence of the exon 11’ showing the start of the cryptic exon (green) 13bp downstream of the mutation (red). The fourth codon of 11’ is a stop codon (underlined).

In OGI-601-1236, the c.1611+27G>A mutation led to a splicing pattern that became apparent only through amplicon sequencing (Figures 3b, S4, S5). In the WT allele, three splicing signals of various strengths were detected between exons 12 and 13 (Figure 3b). The two prominent splice variants have been annotated for *RPGRIP1*, the main event (9598 reads) being present in the NM_020366 transcript and indicating the canonical splice junction between exon 12 and 13. The second splice donor occurring 104bp into intron 12 (8499 reads) is part of the 5’UTR of predicted transcript model XM_005267881.3 (Figure 3b). The third splice donor site occurs 227bp into intron 12 and has not been reported. In both the positive control, c.1611+1G>A, and the mutation under study, c.1611+27G>A, the primary splicing event that would lead to full-length protein failed to occur whereas the alternate transcript donor site, 104 past the NM_020366 exon 12, was preferred (Figure 3b). This variant would lead to a pre-mature stop codon 12 codons after the exon 12 of the full-length transcript (i.e. NM_020366).

In OGI-949-1907, the intronic c.1468-263G>C variant identified resulted in an activation of a cryptic splice acceptor site 13bp past the mutation (Figure 3c). The resultant cryptic exon 11’, although 117bp and in-frame, has a premature stop codon four codons past the cryptic acceptor site. Thus the aberrant transcript will lead to NMD.

In order to account for potential differences in splicing between the neuronal and non-neuronal cell types^29^, all the splicing experiments were repeated with a retinoblastoma cell line WERI-Rb1, derived from photoreceptor cells^30^. The splicing changes resulting from the mutations assessed, led to similar patterns in both HEK293T as well as WERI-Rb1 lines (Figure S5). Overall, we found four novel non-coding mutations, four unreported coding mutations in *RPGRIP1* (Figure 4) and we have corrected the transcript model for *RPGRIP1* in the retina with a new TSS and 5’ exon 8kb upstream of the annotated exon 1 (name exon 1n in this paper).

**Figure 4.**
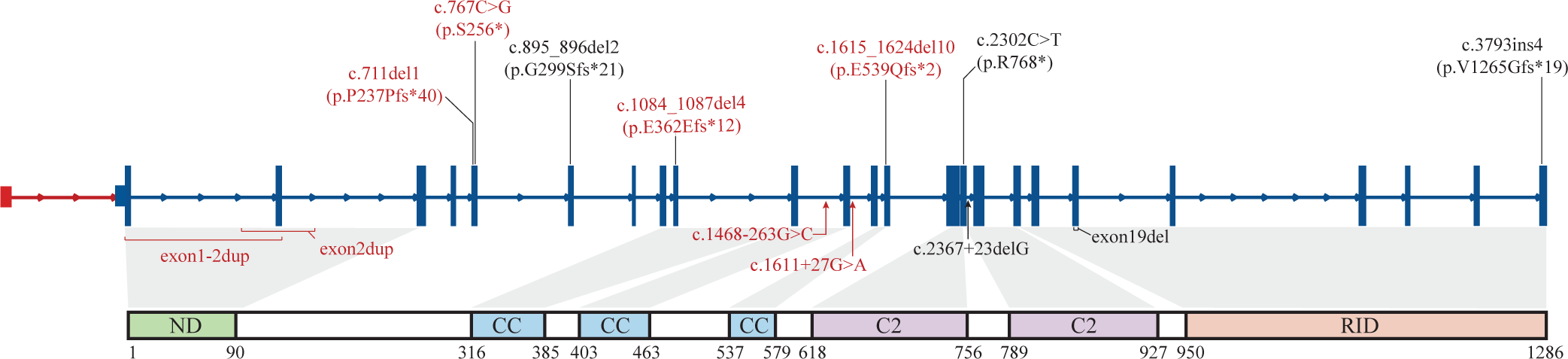
Summary of RPGRIP1 mutations and gene model. Novel findings are indicated in red with non-coding mutations listed below the NM_020366 transcript model and coding mutations listed above. The novel exon 1’ is shown in red. The RPGRIP1 protein and corresponding domains is shown in the bottom.

Among the seven families with one LoF *RPGRIP1* allele, OGI-578 did not reveal a second *RPGRIP1* mutation after WGS and copy number analyses. None of the RPGRIP1 calls showed proper segregation or were of poor quality and failed Sanger validation. WGS however, revelaed other IRD candidate gene, the most prominent of which was *CNGA1* with biallelic high quality calls.

## DISCUSSION

In our analysis of nine families with biallelic *RPGRIP1* mutations, six had a second non-coding mutation leading to a LoF allele. This not only highlights the importance of non-coding mutations in pathogenesis of recessive IRDs, but also implies a greater prevalence of *RPGRIP1* mediated disease than previously thought. Three among the 1722 IRD families had bilallelic coding mutations in *RPGRIP1*, which corresponds roughly to the previously reported rates in the literature (0.17% vs 0.25%^14,31^). However, when taking into account non-coding mutations, our rate increased from 0.17% to 0.51%. This significant increase in frequency of *RPGRIP1* mediated disease is still likely an underestimate given the bias of choosing samples with one LoF coding sequence mutation. Considering the possibility of bialleleic non-coding LoF alleles, the rate of *RPGRIP1* mediated disease is likely to be much greater than what was previously thought.

In the six families solved with the addition of structural and intronic mutation analyses, we validated the second LoF mutations *in vitro* showing aberrations in the reading-frame caused by splicing or copy number changes. While deletions are simpler to interpret given clear interruption of reading frame, copy number gains can be just as disruptive. In both cases of copy number gain, OGI-237 and OGI-281, the mutations were tandem duplications leading to premature stop codons. While investigating the potentially pathogenic effect of the duplication in the first two annotated *RPGRIP1* exons in OGI-237, we discovered a novel exon and a retinal TSS 8Kb upstream of the previously annotated transcriptional start site. This is important as mutations in regulatory regions and 5’UTR have been shown to be disease-causing, such as in the case of *PRPF31* and *OPN1MW^9^*. We propose the inclusion of this *RPGRIP1* exon, chr14:21748265-21748318 (hg19), in future IRD diagnostic panels.

We also detected three mutations causing intron retentions or inclusion of cryptic exons in the resulting transcripts, two of which were in the flanking 30bp of annotated exons. The traditional way of assessing for the effect of intronic mutations includes use of exon-trapping vectors with a small (mini-gene) or medium (midi) sized insert harboring the mutation under study^32,33^. Subsequent transfection into HEK293T cells, and analysis of the processed RNA via RT-PCR allows one to assess the alterations via gel electrophoresis. However, as seen in the case of OGI-601 (Figures 3b, S4b), there can be a multitude of splicing signals, some of which are not annotated for the genes of interest thus making the interpretation of the electrophoresis results difficult. We found that amplicon sequencing of the RT-PCR product can increase the sensitivity of detection (OGI-601, Figures 3b, S4b) and also add a quantitative value to assessment of splicing changes. Additionally, the mutation c.2367+23delG, had been noted previously^3^ as a potential disease causing and splicing altering mutation. Yet its precise effect proved challenging to interpret, while the authors suggested a possible exon 15 and 16 skipping in blood mRNA. Through amplicon sequencing (Figures 3a, S5) we found evidence of intron 15 retention in both HEK293T and the tissue relevant WERI-Rb1 line (Figure S5). Amplicon sequencing analysis clarifies the exact splicing events and proportions thus offering an advantage in interpreting results. It should be noted however, that we cannot rule out exon 15-16 skipping as suggested by Riera *et al.* because exon 17 was not part of the cloned *RPGRIP1* used in the splicing assay. Weakening of splicing between exons 15 and 16 with this mutation is supportive of both models.

Splicing is a tissue-specific phenomenon and neuronal tissues have a unique splicing machinery that can lead to unique exonic retentions^29^. Thus the splicing patterns we identified in HEK293T cells were confirmed in a retina relevant line. We chose WERI-Rb1, a retinoblastoma line shown to possess the neuronal specific splicing machinery^29^. Although we did not detect a significant difference between the two lines (Figure S5), we foresee a benefit in using a tissue relevant cell line, such as WERI-Rb1 for IRDs, when assessing splicing mutations.

Despite the decreasing costs of whole genome sequencing, it is still not feasible for routine clinical assessment. Hence, assessing for non-coding mutations in *RPGRIP1* and other recessive IRD genes still demands a step-wise approach to reduce screening costs. Our recommendation is to include and analyze at least 30bp of the flanking exonic regions and to include copy number analysis^4^ in targeted panel sequencing of recessive IRD genes. In this study, we noted only one significant mutation outside of these criteria (c.1468-263G>C). The remaining unsolved cases can then be submitted for whole genome sequencing to identify novel genes and mutations.

As demonstrated above, the empirical validation of second-mutant alleles in recessive diseases when one mutant allele is identified can be time consuming and expensive. One might argue that such stringency may not be necessary if the gene identified with a single mutation matches the presenting phenotype. Statistical models predict a small false positive diagnosis rate when a single mutation is identified in a disease-specific gene such as *MYO7A* in Usher Syndrome Type I^34^. Yet when a large number of potential genes can lead to the phenotype of interest, such as in early-onset IRD, the calculations will not yield as small a false positive rate^34^. Thus we strongly believe that in order to make the correct diagnosis, it is important to identify and characterize both mutant alleles.

Accurate genetic diagnostics for inherited retinal degenerations is increasingly critical with the emergence of gene therapy. Clinical trials of gene augmentation therapy for *RPE65*-associated retinal degeneration have been completed, and the treatment was recently approved by the FDA^6^. Further, clinical trials of gene therapies for eight other genetic forms of retinal degeneration are in progress^35–40^. Successful pre-clinical studies of gene therapies for multiple other genetic forms of IRD have been reported, including for *RPGRIP1*-associated IRD^7,8^. Evidence of biallelic gene mutations are critical for inclusion of patients in trials when dealing with autosomal recessive diseases. We believe the results reported here are also relevant for other genetic forms of IRD, and that genetic testing which incorporates detection of non-coding and structural mutations will increase the diagnostic sensitivity for all IRDs.

## Acknowledgments

This work was supported by grants from the National Eye Institute [RO1EY012910 (EAP), R01EY026904 (KMB/EAP) and P30EY014104 (MEEI core support)], and the Foundation Fighting Blindness (USA, EAP). Sequencing and analysis was provided by the Center for Mendelian Genomics at the Broad Institute of MIT and Harvard and was funded by the National Human Genome Research Institute, the National Eye Institute, and the National Heart, Lung and Blood Institute grant UM1 HG008900 to Daniel MacArthur and Heidi Rehm. The authors would like to thank the patients and their family members for their participation in this study and the Ocular Genomics Institute Genomics Core members for their experimental assistance. The authors would like to thank the Exome Aggregation Consortium, the Genome Aggregation Database (gnomAD) and the groups that provided exome variant data for comparison. A full list of contributing groups can be found at http://exac.broadinstitute.org/about and http://gnomad.broadinstitute.org/about.

## Supplementary figure legends

***Figure S1.* Overview of the approach.** Of the 1722 families that underwent GEDi sequencing, ten had at least one loss of function (LoF) mutation in *RPGRIP1.* This included three families that were solved with biallelic LoF mutations and seven families that underwent further investigation. One family, OGI-578, failed validation.

***Figure S2.* Validation of exon 19 deletion in pedigree OGI-1797. (a)** The pedigree of the family. **(b)** qPCR of the proband OGI-1797-3128 showing deletion of exon 19 in one allele. The NM_020366 transcript model is shown at the bottom.

***Figure S3.* Confirmation of tandem duplication in OGI-237. (a)** the model of the mutant allele (M4) with the breakpoint of tandem duplication indicated via black arrowhead. The approximate corresponding locations of the primers used in figure S3b are indicated with red, blue (breakpoint PCR) and black (control PCR) arrows. **(b)** PCR confirmation of the tandem duplication using primers across the breakpoint. HEK293T was used as negative control as the mutation was expected to be missing in this cell line. Primers at a distant *RPGRIP1* locus (exon 14-15) were used to control for PCR reaction and DNA quality.

***Figure S4.* Midi-gene splicing assay of intronic mutation.** Gel images of the RT-PCR reactions are shown to the right of each pedigree. Two different cell lines, WERI-Rb1 and HEK293T, were transfected with vectors containing WT, mutant and generated positive control mutants of the *RPGRIP1* mutations seen in families OGI-949 **(a)**, OGI-601 **(b)**, and OGI-827**(c)**. These PCR products were used for the subsequent amplicon-seq analyses.

***Figure S5.* Amplicon-sequencing showing similar splicing pattern in HEK293T and WERI-Rb1 lines.** Minimum split read counts of 1500-5000 were used. The results show lack of significant difference between the different cell lines and confirm the results reported in Figure 3.

***Table S1.* List of primers used and their indications.** All are listed in 5’ to 3’ directionality.

